# N4-acetylcytidine (ac4C) promotes mRNA localization to stress granules

**DOI:** 10.1101/2023.05.22.541693

**Authors:** Pavel Kudrin, Ankita Singh, David Meierhofer, Anna Kuśnierczyk, Ulf Andersson Vang Ørom

**Author notes:** Lead Author, correspondence to (UAVØ).

## Abstract

Stress granules are an integral part of the stress response that are formed from non-translating mRNAs aggregated with proteins. While much is known about stress granules, the factors that drive their mRNA localization are incompletely described. Modification of mRNA can alter the properties of the nucleobases and affect processes as translation, splicing and localization of individual transcript. Here, we show that the RNA modification N4-acetylcytidine (ac4C) on mRNA associates with transcripts enriched in stress granules and that stress granule localized transcripts with ac4C are particularly translationally regulated. In addition, we show that ac4C on mRNA can mediate co-localization of the protein NOP58 to stress granules. Our results show that acetylation of mRNA regulates localization of both stress-sensitive transcripts and RNA-binding proteins to stress granules and adds to our understanding of the molecular mechanisms responsible for stress granule formation.

## Introduction

Stress granules (SG) are membrane-less organelles formed by protein-RNA aggregates upon stress, that are evolutionary conserved across eukaryotes ^1, 2^. SGs have been extensively studied, and while it is well established that they form when translation initiation is limited ^1^ and a variety of roles for SG within the cell has been proposed, their formation, dispersal and function remain largely unclear ^2^.

RNA modifications occur at all RNA species, particularly at rRNA and tRNA, but also mRNA is increasingly reported to contain modified nucleosides ^3^. Particularly N6-methyladenosine (m6A) has been shown to play important roles for mRNA translation ^4^. And while m6A has also been proposed to play a role in mRNA localization to SG through interaction with YTHDF ^5^, a functional relationship has recently been questioned ^6^. More recently, the RNA modification N4-acetylcytidine (ac4C) has been shown to be deposited on mRNA and regulate translation efficiency ^7–9^. ac4C is conserved through all kingdoms of life and is induced upon several different stresses ^7^. ac4C is less abundant than m6A on mRNA and due to difficulties in precise and quantitative mapping its function and occurrence on mRNA has remained controversial ^7, 8^. A recent study using nucleotide-resolution mapping of ac4C in HeLa cells identifies more than 6,000 acetylation sites in mRNA, demonstrating a wide-spread occurrence of acetylation on human mRNA ^9^.

Here, we show that ac4C is enriched in SG and that acetylated transcripts are predominantly localized to SG upon arsenite stress. We propose that acetylation of RNA can drive both mRNA and RNA-binding protein localization to SG, providing new insight into both the function of mRNA acetylation and mechanism of RNA localization to SG.

## Results

### Acetylated RNA is enriched in membrane-less organelles

ac4C is known to occur on 18S rRNA that is highly present in the nucleolus ^10^, where the acetyltransferase NAT10 is also predominantly localized ^11^. In addition, ac4C is induced by oxidative stress from archaea to mammals ^7^. To assess ac4C distribution in the cell during non-stressed and stressed conditions we used microscopy and staining for nucleolar marker NCL and the SG marker G3BP (Figure 1a-e). Here, we used WT HeLa cells and a HeLa cell line where the ac4C acetyltransferase NAT10 has been inactivated (NAT10 KO) ^8^, where the ac4C levels are reduced by 80 per cent on all RNA species.

**Figure 1.**
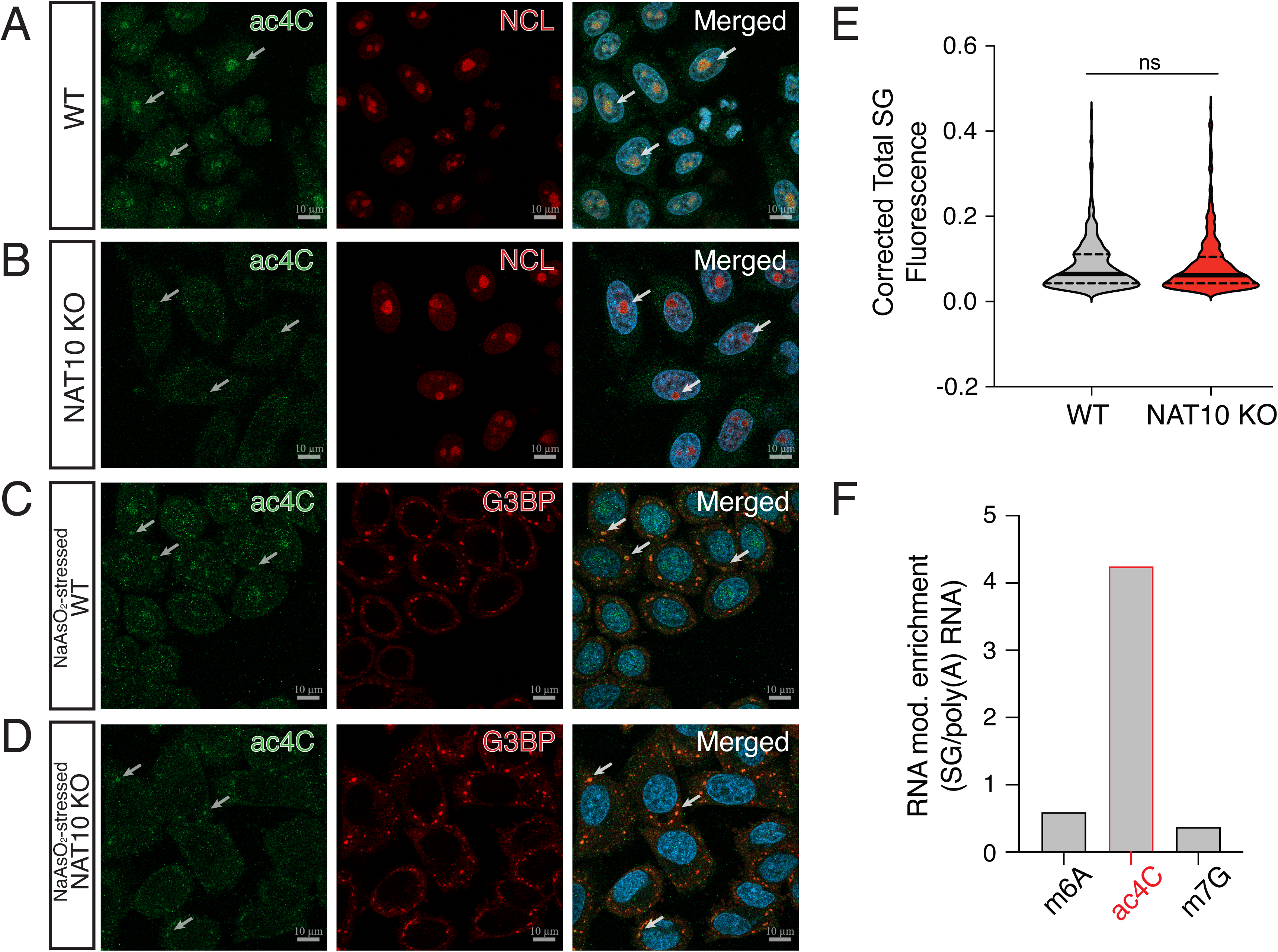
ac4C enrichment in stress granules. Panel a-d show confocal microscopy images of WT (a) and NAT10 KO HeLa cells (b) stained for ac4C and the nucleolus marker Nucleolin and arsenite stressed WT (c) and NAT10 KO HeLa cells (d) stained for ac4C and the SG marker G3BP. Arrows indicate ac4C granules overlapping with nucleoli in (a-b) and SG in (c-d). In (e) the intensities of SG from (c) and (d) are quantified and shown as corrected total SG fluorescence. Median (solid) and quartiles (dashed), unpaired two-tailed Student’s t-test. In (f) is shown the ratio between SG levels of m6A, ac4C and m7G compared to their respective levels on mRNA.

In unstressed conditions we see that ac4C localize to nucleoli both in WT and NAT10 KO cells (Figure 1a-b), albeit with different intensities due to the different baseline levels of ac4C. Upon oxidative stress caused by arsenite we see formation of SG in both WT and NAT10 KO cells as visualized by staining for G3BP. We see that ac4C co-localize with the SG marker in both WT and NAT10 KO cells (Figure 1c-d), demonstrating a shuttling of acetylated RNA. Depletion of NAT10 did not affect the formation nor the size of SG in our experiments, determined by intensity of G3BP staining (Figure 1e), suggesting that ac4C is involved in but not required for SG formation.

As the high concentration of mRNA in SG could lead to the increased ac4C signal seen with immunofluorescence, we purified RNA from SG as described ^1^ and performed RNA mass spectrometry. In addition, we purified poly(A) RNA, total RNA, 18S rRNA and a tRNA fraction (Supplementary Figure 1) for comparison and contamination control. We used the common mRNA modifications m6A known to be deposited at mRNA and proposed to be involved in mRNA recruitment to SG ^5^ and m7G that is present as a 5’cap at mRNAs and also at relatively high levels at tRNA (Supplementary Figure 1). In addition, m6A and m7G are present at different ratios at 18S rRNA and tRNA (Supplementary Figure 1d, e), making contamination with these RNA species in SG and poly(A) RNA purifications possible. When we compare SG levels of m6A, ac4C and m7G to their levels in the mRNA purification, we see a 4.2-fold enrichment of ac4C in SG whereas m6A and m7G show relative levels of 0.59 and 0.37 fold, respectively (Figure 1f). This shows that, while an enrichment is not reflected in the common mRNA modifications m6A and m7G, we show an enrichment of ac4C supporting the observations from immunofluorescence that ac4C modified RNA are indeed enriched in SG. Especially the lack of enrichment of m7G, present at relatively high levels in 18S rRNA (Supplementary Figure 1d), argues against a contamination with rRNA as the reason for increased ac4C in SG.

### Stress granule purification in HeLa cells is highly reproducible

To address the impact of ac4C on the SG transcriptome, we sequenced total RNA from SG with random primers and without rRNA depletion to obtain the complete picture of the SG RNA content. We sequenced total mRNA (including long ncRNAs) from arsenite stressed cells using random primers after rRNA depletion for comparison to SG to determine relative transcript enrichment (Figure 2a-b, Supplementary Table 1). To assess the reproducibility of the protocol and evaluate the quality of our SG purifications, we compared to two previous studies using the same protocol in U2OS cells ^1, 12^. One study used the G3BP as marker for SG purification ^1^, while the other used the SG marker PABP ^12^. We find a substantial and significant overlap of enriched (Figure 2c) and depleted (Figure 2d) transcripts in SG, especially the different cell type taken into account. For this initial comparison we used a cut-off of 2-fold enrichment and p<0.05. We considered the transcripts overlapping in all three datasets as high confidence and compared translation efficiency and mRNA length ^1^ for transcripts unique to HeLa cells between these studies and the high confidence set. For the translation efficiency we do not see a significant difference between the two sets (Figure 2e), whereas for transcript length (Figure 2f) the mRNAs in the high confidence set are significantly longer on average. As transcript length is one of the well-described properties directing mRNA to SG, we increased the fold change cut-off to >4-fold to favor inclusion of high confidence transcripts for the following analysis.

**Figure 2.**
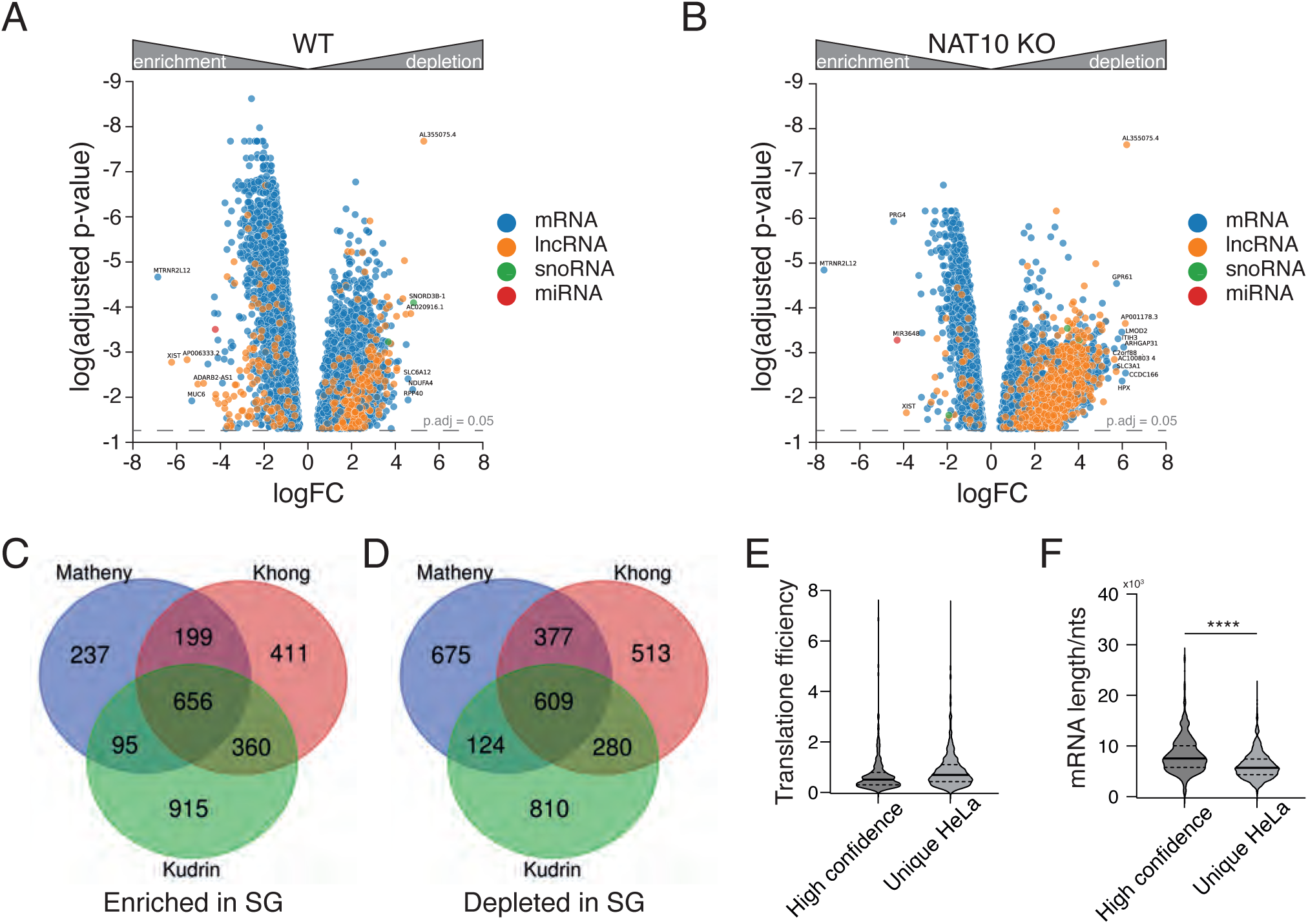
Purification of stress granule-associated RNA. Panel a-b show volcano plots of SG compared to total RNA for WT (a) and NAT10 KO HeLa cells (b) from RNA sequencing experiments done in four biological replicates. We compared enriched (c) and depleted (d) transcripts with previous studies purifying SG from U2OS cells ^1, 12^. We assessed translation efficiency (e) and mRNA length (f) for high-confidence SG transcripts enriched >2-fold in all studies and those enriched only in HeLa cells in the present study, respectively. Unpaired two-tailed Student’s t-test,**** p < 0.0001.

### The stress granule transcriptome is acetylated

Comparing transcript levels in SG to total RNA, we find 418 transcript that are enriched more than 4 times in SG compared to total RNA in WT cells (with a p.adj.<0.05), whereas in NAT10 KO cells only 55 transcripts are enriched using these same criteria, suggesting that the SG RNA content in NAT10 KO cells is less pronounced and closer to the average distribution of mRNA in the cell. When compared to acetylation status using data from [9] we find that 223 of the 418 (53 per cent) SG enriched transcripts in WT cells are acetylated (Figure 3a), which is a significantly higher fraction than expected.

**Figure 3.**
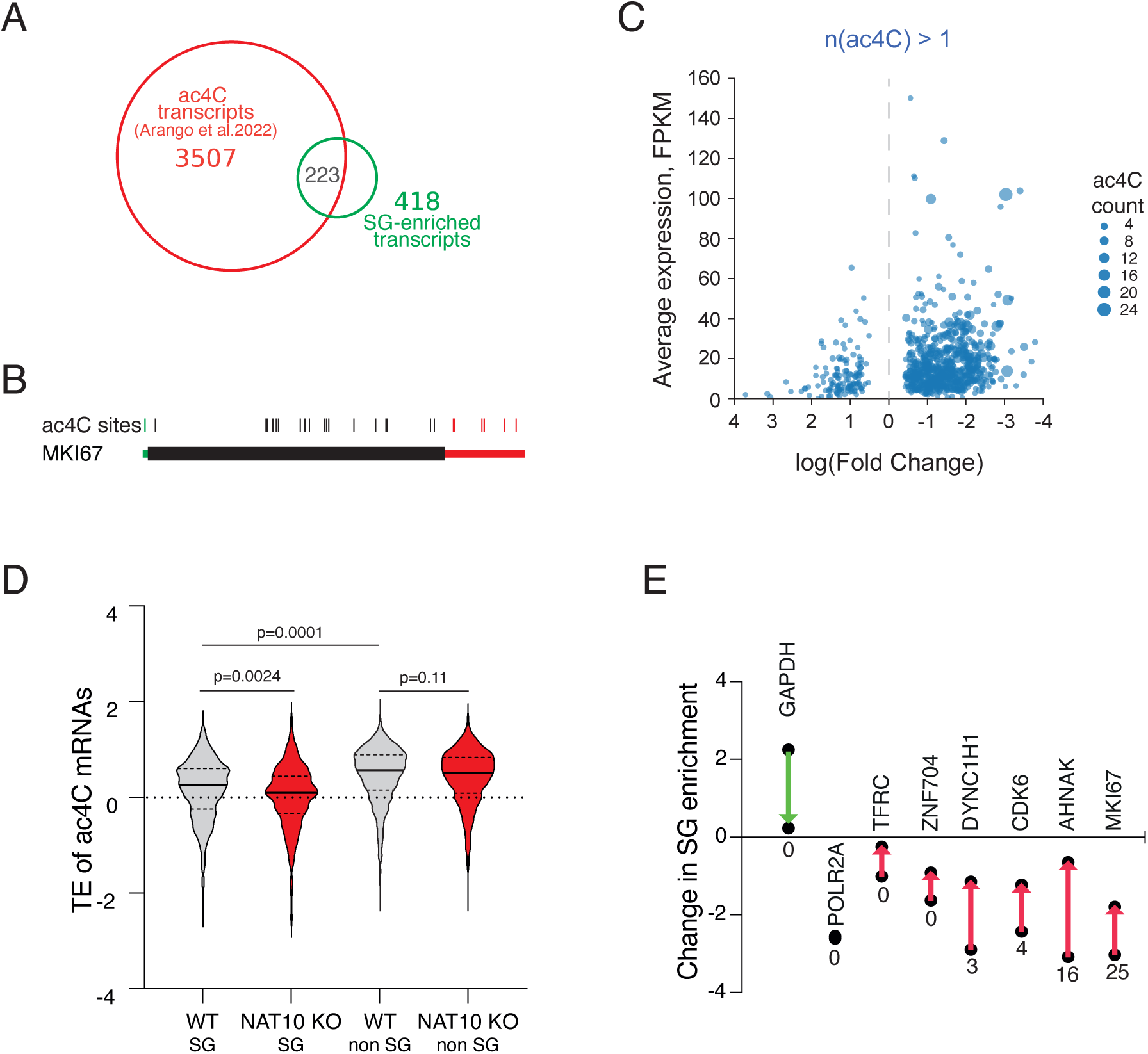
The ac4C-dependent stress granule transcriptome. The overlap between acetylated transcripts in WT HeLa cells and transcripts enriched in SG (53.3 per cent) is shown as a Venn diagram in panel (a). The transcript with the most ac4C sites, MKI67, is shown as a schematic in panel (b), where the 5’ UTR is green, the CDS with the major part of ac4C sites is black and the 3’ UTR is red. Panel (c) shows the transcripts with more than one ac4C sites along with fold change enrichment in SG and normalized average expression shown as FPKM. For the further analysis we used a cut-off of four-fold enriched in SG, and comparison of TE for this set is shown in panel (d) where SG transcripts are defined as having a logFC <-2, and transcripts not enriched in SG are defined as having a logFC >-1. In (e) are shown 8 model transcripts used for comprehensive SG studies ^1^ and their change in SG localization from WT to NAT10 cells. The arrow indicates the direction of the change and below each data point is shown the number of acetylation sites for each transcript, respectively.

Most acetylated mRNAs in the transcriptome are found to have a single site for ac4C modification (56.6 per cent), while a subset display several acetylation sites, up to 25 for the MKI67 transcript ^9^ (Figure 3b). 8 transcripts contain more than 15 ac4C sites, and 7 of these are enriched more than 4-fold in WT SG, reflected in the general observation that ac4C modified transcripts tend to localize to SG upon arsenite stress (Figure 3c and Supplementary Figure 2a-d). The average number of ac4C sites in acetylated SG transcripts is 3.7 compared to 2.0 for the average acetylated transcript in HeLa cells, showing that mRNAs with several ac4C sites are more likely to be localized to SG.

### Translation efficiency is dependent on ac4C level

The mRNAs with high number of ac4C sites have low translation efficiency (TE) (using ribosome profiling data for WT and NAT10 KO cells from ^8^). This is according to the observation that SG are composed of long and less efficiently translated transcripts ^1^, that often correlate with lower TE determined by ribosome profiling ^13^. When looking at TE of acetylated transcripts enriched in SG compared to those that are less than 2-fold enriched, the ac4C transcripts localized to SG (enriched more than four times, logFC<-2) have lower TE on average than acetylated transcripts not enriched in SG (logFC>-1) (Figure 3d). Interestingly, in the NAT10 KO cells with depletion of ac4C, the ac4C transcripts enriched in SG have a lower TE on average than in WT cells, while TE for ac4C transcripts not enriched in SG are not affected on average, suggesting that translation of transcripts that are prone to accumulate in SG are particularly sensitive to acetylation status.

When we look at ac4C status in HeLa cells of the model transcripts used by ^1^, we see that the non-SG control GAPDH is not acetylated. The intermediately SG enriched transcripts POL2RA and TFRC show no acetylation as well. For the SG enriched transcripts DYNC1H1, ZNF704, CDK6 and AHNAK all but ZNF704 are acetylated, as well as the top acetylated transcript MKI67 highly enriched in SG (Figure 3e). The enrichment in SG compared to total RNA of all model transcripts changes between WT cells and NAT10 KO cells towards less pronounced, i.e. SG and total RNA composition becomes more similar in the absence of ac4C (Figure 3e). We observe, that those transcripts that are acetylated undergo the largest change in SG enrichment from WT cells to NAT10 KO cells, whereas e.g. POLR2A that in our data is highly SG enriched, does not change its distribution in the NAT10 KO cells compared to WT cells.

### RNA localization detected with smFISH

To further substantiate the localization of acetylated RNA to SG we used smFISH to visualize the control non-SG mRNA GAPDH (Figure 4a) and the core SG mRNA AHNAK (Figure4b) localization to SG following arsenite treatment.

**Figure 4.**
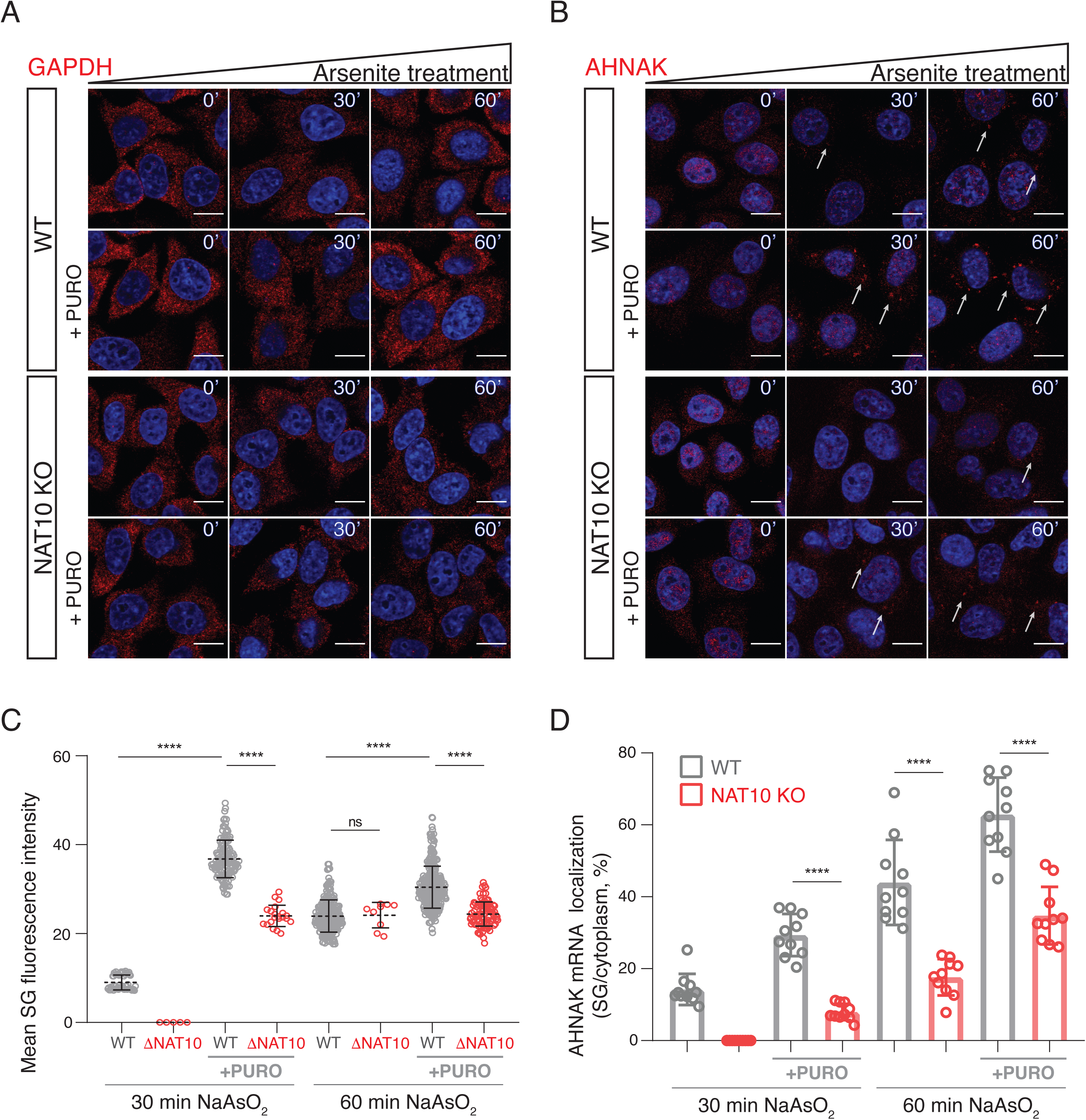
smFISH analysis of AHNAK and GAPDH mRNA localization to SG. We used smFISH to further substantiate our findings using the core SG transcript AHNAK and the control transcript GAPDH. In (a) is shown GAPDH localization and in (b) AHNAK localization upon arsenite stress and puromycin treatment. Arrows indicate AHNAK granules, formed in cytoplasm in response to arsenite stress w/o puromycin treatment. Red is smFISH probe and blue is DAPI. Scale bar, 50 μm. Number and intensity of smFISH signal in individual SGs is quantified from 60 cells for AHNAK in panel (c). Each circle represents a single SG. Unpaired two-tailed Student’s t-test, **** p < 0.0001. Total SG fluorescence per cell, calculated from smFISH signal for AHNAK of individual SGs (c), is shown in panel (d).

ac4C promotes mRNA decoding efficiency ^8^, and it is thus possible that long mRNAs with ac4C modifications exit translation less efficiently in ac4C depleted NAT10 KO cells. This would result in mRNAs remaining in polysomes and not partitioning into SG. To assess if this is the case we used puromycin treatment at the same time as arsenite treatment (shown in Figure 4a,b), where puromycin releases residual ribosomes that would inhibit partitioning of mRNA into SG ^14^. We observe, that puromycin treatment yields more efficient partitioning of AHNAK into SG in both WT and NAT10 KO conditions with both stronger and more SGs formed (Figure 4c), but that the partitioning into SG is still significantly decreased in NAT10 KO cells. The overall distribution of AHNAK into SGs shows a markedly reduced partitioning into SG in NAT10 KO cells compared to WT (Figure 4d), and while the puromycin effect is clearly promoting SG formation it does not overrule the impact of ac4C modification of partitioned transcripts. The expression on mRNA level of AHNAK is higher in NAT10 KO cells by RNA sequencing, further supporting that the decreases partitioning of AHNAK into SG in NAT10 KO cells is an effect of ac4C rather than of mRNA level in the cell. Thus our data suggest that ac4C directly affect the targeting of RNA into SG rather than indirectly through its reported effect on decoding efficiency ^8^.

### Identification of ac4C binding proteins

To identify protein binders recognizing the ac4C modification, we in vitro synthesized three biotinylated 76 nts RNAs containing 21, 24 and 16 C nucleotides, respectively, in various sequence contexts. The RNA was synthesized in the presence of unlabeled A, U and G ribonucleotides and increasing concentrations of ac4C ribonucleotides (0, 50 and 100 per cent ac4C over C, respectively). The in vitro synthesized RNA was incubated with total cellular lysate from NAT10 KO cells, to increase the unbound fraction of ac4C binders. We eluted bound proteins from RNA using excess biotin and RNase, and subjected purified proteins to liquid-phase mass spectrometry (LC/MS). Here, we identify 18 proteins preferentially bound by the ac4C labeled RNA compared to the control non-acetylated RNA (Figure 5a, Supplementary Figure 4, Supplementary Table 2).

**Figure 5.**
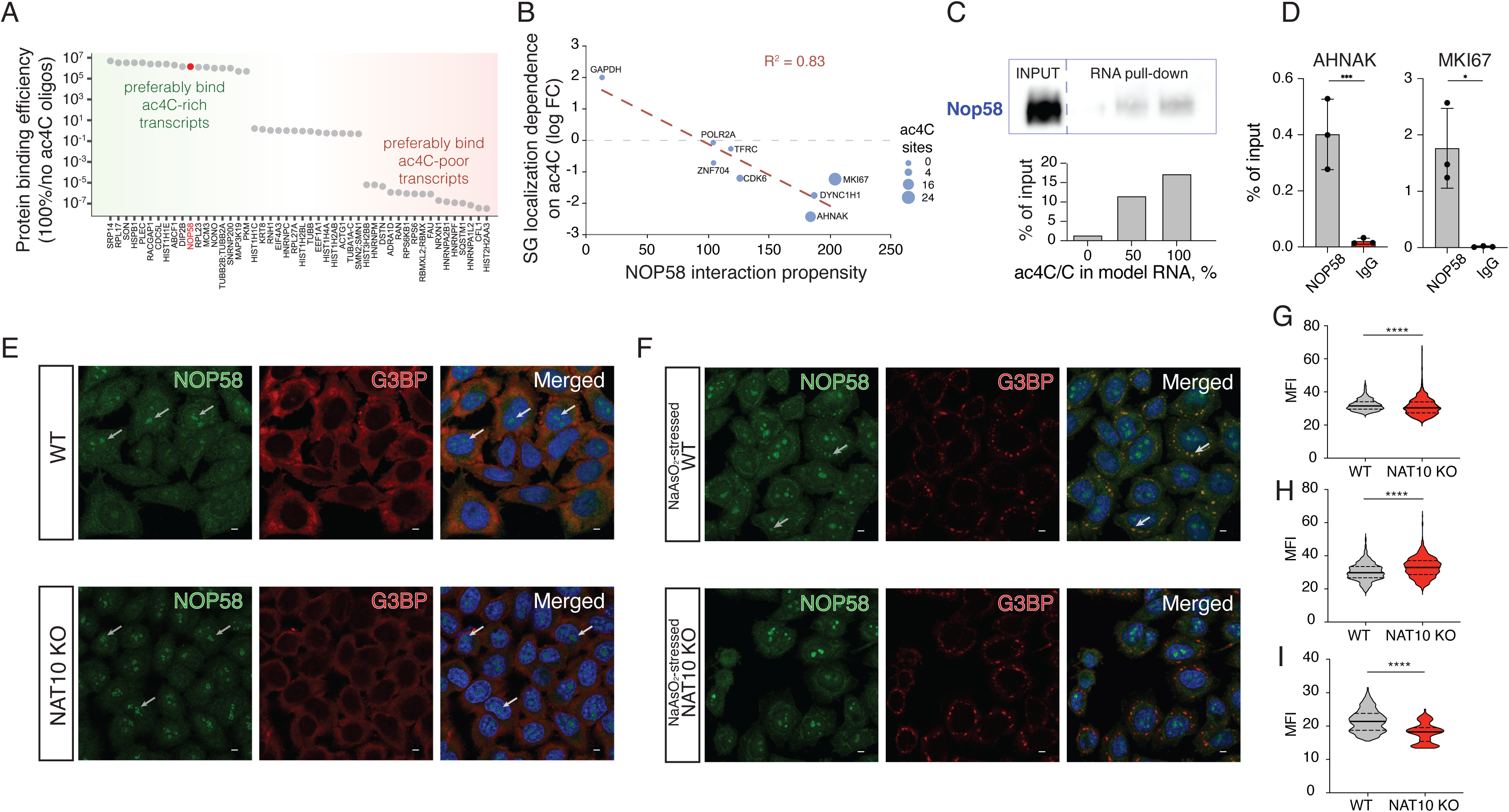
NOP58 localization to SG is dependent on ac4C. Panel a shows proteins identified to bind preferentially to ac4C modified RNA oligonucleotides as well as those binding regardless of acetylation status and preferentially to unacetylated RNA oligonucleotides. (b) The change in SG enrichment from WT to NAT10 KO HeLa cells for the SG model transcripts is shown as a function of interaction propensity with NOP58 predicted by Catrapid. The size of the datapoints show the number of acetylation sites on each transcript. (c) Dose-dependent interaction of NOP58 to acetylated RNA oligonucleotides validated by western blot (upper panel) and quantified as compared to 100 per cent input (lower panel). (d) RIP-qPCR of NOP58 interaction with the SG core transcript AHNAK as well as for MKI67 in WT HeLa cells. Immunostaining of NOP58 and G3BP for unstressed cells are shown in panel (e) and for arsenite stressed cells shown in panel (f). Arrows indicate NOP58 granules overlapping with nucleoli in (e) and SG in (f). NOP58 containing SG are not indicated in NAT10 KO HeLa cells in (f) as they are not visible. Scale bar, 10 μm. Panel (g-h) show quantification of NOP58 Mean Fluorescence Intensity (MFI) in nucleoli of unstressed (g) and arsenite stressed cells (h). In (i) is shown quantification of MFI of SG in arsenite stressed WT and NAT10 KO HeLa cells. Unpaired two-tailed Student’s t-test, **** p<0.0001.

Due to the artificial nature of the acetylated RNA oligonucleotides and as a complementary approach to identify biological relevant proteins interacting with the SG enriched transcripts, we used Catrapid ^15^ to predict proteins interacting with MKI67, that is highly enriched in SG and contains the most ac4C sites of all transcripts in HeLa. Here, we find that NOP58 is the top predicted protein to bind MKI67, which is also one of the best hits of proteins identified to bind acetylated RNA in our mass spectrometry analysis. Overall, when using the model transcripts GAPDH, POL2RA, TFRC, DYNC1H1, ZNF704, CDK6, AHNAK and MKI67 we see a high degree of correlation between the interaction propensity score from Catrapid as a function of change in SG localization between WT cells and NAT10 KO cells (Figure 5b), supporting the identification of NOP58 as a bona fide ac4C binder.

Due to the experimental identification and prediction of NOP58 to bind to acetylated SG transcripts we focused our further validation and functional characterization of ac4C binding proteins in SG localization on NOP58. NOP58 is a nucleolar protein that has also been shown to be localized to SG ^16^. In unstressed cells, ac4C is highly present in the nucleolus. We validate the interaction between ac4C and NOP58 by repeating the pull-down procedure and western blotting using a NOP58-specific antibody, confirming a dose-dependent binding between ac4C on RNA and NOP58 (Figure 5c), and using RIP-qPCR, compared to IgG control, we further validated the interaction between NOP58 and the core SG mRNA AHNAK, having 16 ac4C sites, and the most acetylated mRNA MKI67, having 25 ac4C sites (Figure 5d).

### NOP58 is binding to acetylated mRNA and localized to stress granules

To study the cellular connection between ac4C and NOP58 localization to SG we stained for NOP58 in WT and NAT10 KO cells either in unstressed cells or in cells treated with arsenite. In untreated cells, NOP58 localizes to nucleoli in both WT and NAT10 KO HeLa cells (Figure 5e). Upon arsenite stress and induction of SG we see that in WT HeLa cells NOP58 is recruited to SG (Figure 5f, upper panel) (in agreement with the SG proteome from ^16^), whereas in NAT10 KO this recruitment is abrogated (Figure 5f, lower panel). Quantification of NOP58 levels show that NAT10 KO HeLa cells have higher levels of NOP58 in the nucleoli (Figure 5g-h), and that SG levels of NOP58 are significantly higher in WT compared to NAT10 KO HeLa cells. This suggests that ac4C modified RNA binds NOP58 (and other proteins) and is able to recruit them to SG, proposing that acetylated RNA is important for shaping both RNA and protein content of SG.

## Discussion

A recent review on SG asked the question, how does RNA contribute to SG formation, and which RNAs are important? ^2^. Several studies using diverse approaches have suggested that transcript length is the key determinant of mRNA recruitment to SG ^1, 12, 17^. Here, we show that transcripts modified with ac4C are particularly important for defining the diversity of mRNA in SG. The formation of SG is maintained in NAT10 KO HeLa cells albeit with different mRNA content, showing that SG can still form but the mRNA distribution is more similar to the average mRNA distribution of the cell. We do not see relative enrichment of m6A in SG which could be expected due to the observation that m6A mRNA is transported to SG by YTHDF ^5^, but this lack of enrichment of m6A is in line with a recent study showing a modest impact of m6A on SG RNA composition ^6^.

Our findings that ac4C affects the composition and distinctness of SG mRNA content adds the question how diverse SG content are across cell lines, tissues and external stimuli, and how this is affected by different acetylation patterns across cell lines and tissues. Part of this question might be answered more in-depth once we have a more comprehensive picture of RNA ac4C in a panel of cell lines and tissues, but our comparison with previous SG purification from U2OS cells identifying high-confidence SG transcripts suggest that ac4C levels are partially comparable between cell lines and involved in partitioning of RNA into SG.

Our data also suggest that mRNA is not passively dragged along to SG but are localized there dependent on acetylation status and can mediate protein localization to SG. Why acetylated transcripts that are localized to SG have lower TE and are more susceptible to NAT10 KO than other acetylated transcripts is an outstanding question. It might be associated to the translational status of the transcripts, and possibly indicate that low TE mediate acetylation of transcripts with a subsequent acetylation-dependent enhancement of translation, fitting well with very recent data that ac4C promote translation ^9^. Using puromycin treatment and smFISH we address whether the accumulation of ac4C RNA into SG is due to the rate at which mRNAs exit translation. Here, we see that while puromycin treatment increases the rate of mRNA partitioning into SG it does not override the effect of NAT10 KO mediated decrease in ac4C levels, in support of a model where the presence of ac4C directly affects the partitioning rate of mRNA into SG upon arsenite stress. Our assessment of the ac4C-binding protein NOP58 and its localization into SG also supports a model where the RNA acetylation and partitioning into SG mediates protein localization, and not vice versa, proposing RNA acetylation as an important factor for defining both the transcriptome and proteome of SG induced by arsenite stress.

SGs share many protein components with neuronal granules, and mutations associated with SG formation have been shown to be implicated in neurodegenerative diseases such as amyotrophic lateral sclerosis (ALS) and multisystem proteinopathy, where SG-like assemblies form ^18^. Addressing the acetylation status of RNA could provide novel insight into such diseases.

### Limitations of the study

With the findings presented here we show an involvement of ac4C modification in SG partitioning of RNA. Previous work has shown that particularly long mRNAs tend to be localized to SG, and as the most highly acetylated transcripts are also very long it is not possible from our data to clearly distinguish between the contribution of length and ac4C modifications, respectively. The ac4C interacting proteome and how RNA-protein interactions affect the transcriptome and proteome of SG is of high interest but outside the scope of this article. We hope that our initial data on NOP58 and our findings will foster further investigation of how RNA, RNA modifications and proteins play together to form SG.

Supplementary Information Supplementary Figures 1, 2, 3 and 4. Supplementary Tables 1, 2 and 3.

## Data availability

RNA sequencing data from total RNA and SG have been deposited to GEO with accession number GSE212380. Riboseq data are available from Arango et al., 2018 with accession number GSE102113 and position of ac4C on HeLa mRNA are from Arango et al., 2022 with accession number GSE162043.

## Acknowledgments

We thank Ana Rebane for providing research facilities for PK during parts of the work for the manuscript. We thank Beata Lukaszewska-McGreal for proteome sample preparation. We thank Shalini Oberdoerffer for kindly providing the NAT10 KO and the WT HeLa cell lines. The LC-MS/MS analyses of RNA were performed by the Proteomics and Modomics Experimental Core (PROMEC), Norwegian University of Science and Technology (NTNU) and The Central Norway Regional Health Authority. This facility is a member of the National Network of Advanced Proteomics Infrastructure (NAPI), which is funded by the Research Council of Norway. We acknowledge AU Health Bioimaging Core Facility for the use of equipment and support of the imaging facility. Work in the author’s lab is funded by the Novo Nordisk Foundation, The Lundbeck Foundation, Danish Cancer Society, Independent Research Fund Denmark, The Carlsberg Foundation to UAVØ, Estonian Research Council to PK and the Max Planck Society to DM.

## Author contributions

Designed experiments: PK, UAVØ; Performed Experiments: PK, AK, DM; Performed computational analysis: AS, UAVØ; Interpreted data: PK, AS, AK, DM, UAVØ; Supervised research: UAVØ; Secured funding: UAVØ; Wrote the initial draft: PK, UAVØ; wrote the final manuscript: PK, UAVØ. Commented on and approved the final manuscript: All authors.

## Materials and Methods

### Tissue culture

If not stated otherwise, HeLa cells were grown in Dulbecco’s Modified Eagle’s medium (DMEM; Gibco, #41966-029) supplemented with 10% Fetal Bovine Serum (FBS; Gibco) and 1% Penicillin/Streptomycin (P/S; Gibco) at 37°C with 5% CO_2_ until 90% confluency. Cells were collected by trypsinization with 0.05% Trypsin-EDTA (Gibco). When needed, the purification of SG cores was performed as per ^1^. Briefly, 90% confluent HeLa cells were subjected to oxidative stress by 1 hr treatment with 0.5 mM NaAsO_2_ followed by lysis in SG lysis buffer (50 mM Tris-HCl pH 7.4, 100 mM KOAc, 2 mM MgOAc, 0.5 mM DTT, 50 mg/mL Heparin, 0.5% NP40, 1 mM PMSF, 1:100 PI cocktail, Superase-in (1 U/ml) (Thermo)) and fractionation through centrifugation. Stress granule core enriched fraction was then incubated at 4°C overnight with anti-G3BP1 antibody (#61559, CellSignal). Anti-G3BP1 antibody-bound SG cores were collected through the incubation with Protein G Dynabeads (Invitrogen) with consecutive SG RNA elution from the Dynabeads by Proteinase K (Invitrogen) treatment. Total or SG RNA purification was performed using TRIzol (Invitrogen) according to manufacturer’s instructions and RNA preparation quality analyzed with 2100 Bioanalyzer (Agilent).

### Immunofluorescence staining (IF), RNA FISH and confocal microscopy

HeLa cells were grown on a coverslip and, with 90% confluency reached, were, when needed, subjected to oxidative stress by incubation with 0.5 mM NaAsO_2_ with or without the presence of 10 μg/ml puromycin for either 30 min or 60 min. Cells were fixed with 4% paraformaldehyde in PBS. For IF experiments, permeabilisation and blocking were done with 0.1% Triton-X100 and 0.01% Triton-X100/1% FBS in PBS respectively. Primary antibodies (rabbit anti-ac4C, #ab252215, Abcam; rabbit anti-NOP58, #14409-1-AP, Proteintech; mouse anti-NCL, #87792, CellSignal; mouse anti-G3BP, #ab56574, Abcam) were diluted 1:50 in blocking solution and incubated with the samples for 1 hr at RT. Subsequently the samples were incubated with secondary antibodies (anti-rabbit AlexaFluor488 conjugated and anti-mouse AlexaFluor647 conjugated) at 1:50 dilutions for 1 hr at RT.

RNA FISH experiments were performed according to Stellaris RNA FISH protocol for adherent cells (https://www.biosearchtech.com/support/resources/stellaris-protocols). After fixation with 4% PFA cells were permeabilized by incubation in 70% EtOH for at least 1 hr at 4°C followed by incubation in Stellaris Wash Buffer A (Biosearch Tech) for 5 min at RT. Hybridization step occurred in a parafilm-sealed humidity chamber with cells incubated in 125 nM RNA FISH probes (Supplementary Table 3)-containing Stellaris Hybridization Buffer (Biosearch Tech) for 16 hrs at 37°C in the dark. After hybridization the samples were washed twice with Stellaris Wash Buffer A (Biosearch Tech) for 30 min at 37°C in the dark. Additional washing step with Stellaris Wash Buffer B (Biosearch Tech) was performed for 5 min at RT before mounting the samples on the glass slides.

Samples were mounted on glass slides using SlowFade Gold Antifade Mountant with DAPI (Invitrogen). Confocal images were captured by a Zeiss LSM 800 Airyscan laser scanning microscope or Olympus FV1200MPE multiphoton laser scanning microscope. Zen 2010 or Olympus FluoView FV1000 acquisition software, respectively, and ImageJ (Fiji) were used for imaging and analysis.

### RNA sequencing and data analysis

Total RNA and SG RNA was sequenced with BGI DNBSEQ sequencing technology (BGI). For total RNA samples were depleted for rRNA and library generated with random hexamers. For SG, we did not deplete rRNA to maintain the complete picture of SG composition, and generated libraries with random hexamers.

Coverage of the sequenced libraries was ∼50 million reads (Q20% > 97.2).

### Quality control

Quality control of all of the fastq files were performed with the help of multiqc ^19^. Percentage of uniquely mapped reads was in the range of ∼91-95%, with ∼3-7% of multi-mapped reads.

### Alignment of reads

Alignment of all the paired end reads with the human genome assembly hg19 was performed using STAR (version 2.7.3a) ^20^, Samtools view (version 1.3.1) ^20, 21^ command was used to convert bam files to sam files. Further, quantification of the aligned transcripts from the reference hg19 was performed with the help of htseq-count ^22^ using intersection strict as a mode and stranded yes as the parameters.

### Quantification, differential analysis and annotation

For analysis of RNA expression, readcounts from input samples were used applying a CPM cutoff of 1 or above in all four biological replicates to discriminate expressed genes for the entire dataset. Genes were normalized using the TMM algorithm ^23^ and calcNormFactor of edgeR ^24^. We next used the relationship function voom ^25^ from the limma (Smyth, n.d.) package to establish the mean variance relationship and generate weights for each observation. The lmFit function of limma was used to transform the RNA-Seq data before linear modeling and find differentially expressed (DE) genes.

### In vitro transcription of ac4C modified RNA

T7 promoter containing double stranded DNA (Supplementary Table 3) was used as a template for in vitro transcription with HighYield T7 mRNA Synthesis Kit (ac4CTP) (Jena Bioscience).

DNA template variant 1: sense strand 5’

GTACGGTAATACGACTCACTATAGGGATTGTGCGTGAGATGCACATTCCTGACCGGTGTCTCTTTCTTGACCGGGCCATCCCACATCCGCCGACGC 3’ and antisense strand 5’ GGCCGCGTCGGCGGATGTGGGATGGCCCGGTCAAGAAAGAGACACCGGTCAGGAATGTGCATCTCACGCACAATCCCTATAGTGAGTCGTATTACC 3’. DNA template variant 2: sense strand 5’ GTACGGTAATACGACTCACTATAGGGAGTGGTCTACACACATGACAGAATGGGGCAGGTCCGTAATCGGTTGCAGAGCGGTTACCGATCTCATCGC 3’ and antisense strand 5’ GGCCGCGATGAGATCGGTAACCGCTCTGCAACCGATTACGGACCTGCCCCATTCTGTCATGTGTGTAGACCACTCCCTATAGTGAGTCGTATTACC 3’. DNA template variant 3: sense strand 5’ GTACGGTAATACGACTCACTATAGGGCTTATCTAGTGCATCCGCCGAAATTACCTGTTGCACGACCACGCTCTGCCGCCTCTCAGACTCCTAACGC 3’ and antisense strand 5’ GGCCGCGTTAGGAGTCTGAGAGGCGGCAGAGCGTGGTCGTGCAACAGGTAATTTCGGCGGATGCACTAGATAAGCCCTATAGTGAGTCGTATTACC 3’. Either CTP or ac4CTP substrates were used to obtain the certain level of acetylated cytidines within RNA. Purified RNA was subjected for 3’ end biotinylation with Pierce™ RNA 3’ End Biotinylation Kit (Thermo) resulting in 76 nt long RNA of the following sequences: Variant 1: 5’ GGCCGCGUCGGCGGAUGUGGGAUGGCCCGGUCAAGAAAGAGACACCGGUCAGGAAUGUGCAUCUCACGCAC `AAUC-C`(biotine) 3’ with either 0% or 100% of Cs acetylated. Variant 2: 5’ GGCCGCGAUGAGAUCGGUAACCGCUCUGCAACCGAUUACGGACCUGCCCCAUUCUGUCAUGUGUGUAGACC `ACUC-C`(biotine) 3’ with either 0% or 100% of Cs acetylated. Variant 3: 5’ GGCCGCGUUAGGAGUCUGAGAGGCGGCAGAGCGUGGUCGUGCAACAGGUAAUUUCGGCGGAUGCACUAGAU `AAGC-C`(biotine) 3’ with either 0% or 100% of Cs acetylated.

### Purification of ac4C binding proteins

50 μL of Neutravidine SpeedBeads (Sigma) beads per reaction were equilibrated in buffer A (20 mM Tris-HCl pH 7.4, 1M NaCl, 1 mM PMSF, PI cocktail, Superase-in (1 U/ml) (Thermo), 1 mM EDTA) followed by addition of 50 pmol of biotinylated model RNA with or without acetylated Cs. After the incubation on rotator at RT for 1 h the beads were washed three times and resuspended in buffer B (20 mM Tris (pH 7.4), 50 mM NaCl, 2 mM MgCl2, 0.1% Tween™-20). During the incubation step the HeLa total cell lysate was prepared by lysing freshly collected HeLa cells in RIPA buffer (Sigma) (1 ml per 15 cm plate) supplemented with 1: 100 protease inhibitor cocktail (Sigma) and 1 mM PMSF on ice for 15 min followed by sonication and another 15 min on ice. Cell debris was removed by centrifugation at 4 °C for 10 min at ≥10000 g. Protein concentration was determined by Bradford assay. 100 μg of HeLa lysate per reaction was mixed with 1x buffer B, 15% glycerol and RNase-free H_2_O with consecutive addition on RNA-beads mix and incubation at 4°C for 60 min with rotation. The beads were washed three times with Wash buffer (20 mM Tris (pH 7.4), 10 mM NaCl, 0.1% Tween™-20, 1 mM PMSF, 1:100 PI cocktail) and RNA-bound proteins eluted in 28 μL of elution buffer (1 mM biotin in Wash Buffer and 2 μL RNase) by incubation shaking at 37°C for 30 min. Eluted proteins were subsequently analyzed by PAGE, Western Blot and MS.

### Proteomics Sample Preparation and LC-MS/MS Instrument Settings

Samples were delivered in 1x PBS, 0.01% SDS, the pH was adjusted to 8.5 by adding a final concentration of 100 mM Tris, followed by denaturing at 95°C for 10 minutes at 1000 rpm. 4 µg protein of each sample was further processed. Reduction of cysteines was carried out by adding 1.1 µl of 0.1 M tris(2-carboxyethyl)phosphine at 37°C for 30 minutes at 800rpm, alkylation of cysteines similarly by adding 2.5 µl of 0.2 M 2-chloroacetamide. Samples were digested by trypsin (enzyme-protein ratio 1:40) at 37°C overnight, desalted and reconstituted in 2% formic acid and 5% acetonitrile in water prior to injection to nano-LC-MS. For each sample, 1 and 3 µg protein were injected. LC-MS/MS was carried out by nanoflow reverse phase liquid chromatography (Dionex Ultimate 3000, Thermo Scientific, Waltham, MA) coupled online to a Q-Exactive HF Orbitrap mass spectrometer (Thermo Scientific, Waltham, MA), as reported previously ^26^. Raw MS data were processed with MaxQuant software v1.6.10.43 ^27^, runs from the same samples were combined and searched against the human UniProtKB with 75,074 entries, released in 05/2020.

### Validation of ac4C binding proteins and western blot

The eluate was subjected to SDS PAGE on Novex Tris-Glycine 4-20% (Invitrogen) gel followed by either silver staining with Pierce Silver Stain Kit (Thermo) or Western blotting against anti-NOP58 (#ab236724, Abcam) and anti-GAPDH (#5174s, CellSignal) antibodies. Western blot was developed using Pierce ECL Western Blotting Substrate (Thermo) and imaged with Amersham Imager 600 (GE Healthcare).

### Size-exclusion chromatography of total RNA

Total RNA was fractionated into tRNA, 18S rRNA and 28S rRNA using two dimensions of size-exclusion chromatography (SEC) carried out on an Agilent HP1200 HPLC system with UV detector and fraction collector. The 1^st^ SEC dimension was performed using a Bio SEC-5 1000 Å, 5 μm, 7.8 x 300 mm column (Agilent Technologies, Foster City, CA) and isocratic elution with 100 mM ammonium acetate (pH 7.0) at 500 μl/min for 40 min at 60°C, collecting three fractions containing 28S rRNA, 18S rRNA, and RNAs below 200 nt (‘small RNAs’), respectively. The fractions were lyophilized and the small RNA fraction was reconstituted in 20 μl of water and subjected to a 2^nd^ dimension of SEC using an AdvanceBio SEC 120 Å, 1.9 μm, 4.6 x 300 mm column (Agilent Technologies, Foster City, CA) and isocratic elution with 100 mM ammonium acetate (pH 7.0) run at 150 μl/min for 40 min at 40°C.

### Analysis of isolated RNA species using LC-MS/MS

RNA was hydrolyzed to ribonucleosides by 20 U benzonase (Santa Cruz Biotech) and 0.2 U nuclease P1 (Sigma-Aldrich, Saint-Louis, MO) in 10 mM ammonium acetate pH 6.0 and 1 mM magnesium chloride at 40 °C for 1 hour, then added ammonium bicarbonate to 50 mM, 0.002 U phosphodiesterase I and 0.1 U alkaline phosphatase (Sigma-Aldrich, Saint-Louis, MO) and incubated further at 37 °C for 1 hour. The hydrolysates were added 3 volumes of acetonitrile and centrifuged (16,000 g, 30 min, 4 °C). The supernatants were lyophilized and dissolved in 50 µl water for LC-MS/MS analysis of modified and unmodified ribonucleosides. Chromatographic separation was performed using an Agilent 1290 Infinity II UHPLC system with an ZORBAX RRHD Eclipse Plus C18 150 x 2.1 mm ID (1.8 μm) column protected with an ZORBAX RRHD Eclipse Plus C18 5 x 2.1 mm ID (1.8 µm) guard column (Agilent Technologies, Foster City, CA). The mobile phase consisted of water and methanol (both added 0.1% formic acid) run at 0.23 ml/min, for modifications starting with 5% methanol for 0.5 min followed by a 2.5 min gradient of 5-15 % methanol, a 3 min gradient of 15-95% methanol and 4 min re-equilibration with 5% methanol. A portion of each sample was diluted for the analysis of unmodified ribonucleosides which was chromatographed isocratically with 20% methanol. Mass spectrometric detection was performed using an Agilent 6495 Triple Quadrupole system with electrospray ionization, monitoring the mass transitions 268.1-136.1 (A), 284.1-152.1 (G), 244.1-112.1 (C), 245.1-113.1 (U), 286.1-154.1 (ac^4^C), 282.1-150.1 (m^6^A) and 298.1-166.1 (m^7^G) in positive ionization mode.

### NOP58 binding RNA immunoprecipitation

40 µl of Protein G Dynabeads (Invitrogen) in IP buffer (150 mM NaCl, 10 mM Tris-HCl, pH 7.5, 0.1% IGEPAL CA-630, 1 mM PMSF, 1:100 PI cocktail, Superase-in (1 U/ml) (Thermo) in nuclease free H_2_O) were tumbled with 5 μg anti-NOP58 antibody (#14409-1-AP, Proteintech) or 5 μg anti-IgG antibody (#30000-0-AP, Proteintech) at 4°C for at least 6 hrs. Upon freshly prepared (in RIPA (Sigma) buffer) HeLa cell lysate addition, lysate-antibody-beads mixture in IP buffer was incubated ON at 4°C with gentle rotation in a final volume of 0.8 mL in protein low-binding tubes. For elution, the beads were resuspended in 1x Proteinase K buffer (100 mM Tris-HCl pH 7.5, 150 mM NaCl, 12.5 mM EDTA, 2% SDS and 120 μg/ml Proteinase K (Invitrogen)) and incubated 1 hr with continuous shaking (1200 rpm) at 37°C. Magnetic separation rack was applied to collect the supernatant. TRIzol/chloroform treatment was applied to supernatant with consecutive centrifugation at >13500 rpm for 15 min at 4°C. Upper phase was collected and 700 μl of RLT buffer and 1400 μl of 100% ethanol were added and mixed thoroughly. The mixture was transferred to an RNeasy MiniElute spin column (QIAGEN) and centrifuged at >12000 rpm at 4°C for 1 min. This step was repeated until all sample was loaded to the column. The spin column membrane was washed with 500 μl of RPE buffer once, then 500 μl of 80% ethanol once and centrifuged at full speed for 5 min at 4°C remove the residual ethanol. RNA was eluted with 14 μl ultrapure H_2_O. RNA concentration was measured using the Qubit RNA HS Assay Kit as per the manufacturer’s instructions

### qPCR and RNA purification

To validate stress granule enriched transcripts from previously purified SG RNA by qPCR, cDNA was synthesized with RevertAid First Strand cDNA Synthesis Kit (Thermo) according to manufacturer’s protocol. Platinum™ SYBR™ Green qPCR SuperMix-UDG (Thermo) and the following primers were used for qPCR:

AHNAK FW 5’TCTTCAGCTCCTGCAGCTCT 3’ AHNAK RV 5’CTCCATCTTCCGACTTCAGC 3’ MKI67 FW 5’AGCCCCAACCAAAAGAAAGT 3’ MKI67 RV 5’TTTGTGCCTTCACTTCCACA 3’

### Statistics

All statistics are done using unpaired two-tailed Student’s T-test.

