## Supplemental data for "N4-acetylcytidine (ac4C) promotes mRNA localization to stress granules"

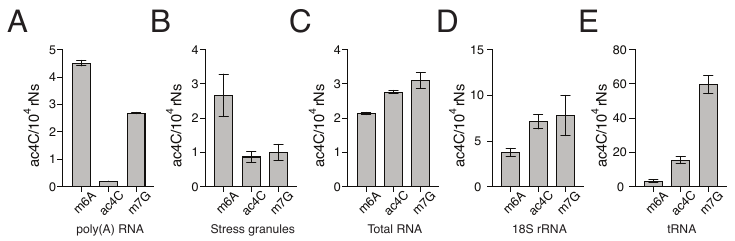

**Supplementary Figure 1. Determination of RNA modification level in SG**

RNA mass spectrometry analysis of m6A, ac4C and m7G levels in (a) poly(A) RNA, (b) SG, (c) total RNA, (d) 18S rRNA, and (c) tRNA from WT HeLa cells. Experiments are done in 2 replicates for panel (a) and three replicates for panels (b-e).

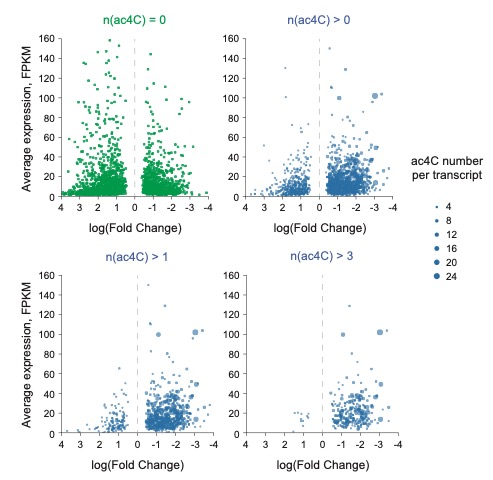

**Supplementary Figure 2. Enrichment in SG, normalized expression and ac4C counts on transcripts**

The figure shows the transcripts with no ac4C sites (a), more than zero ac4C sites (b), more than one ac4C sites (c), and more than three ac4C sites (d) and the found log2FC between SG and total RNA (negative FC means enriched in SG) as well as average normalized expression from RNA-seq shown as FPKM.

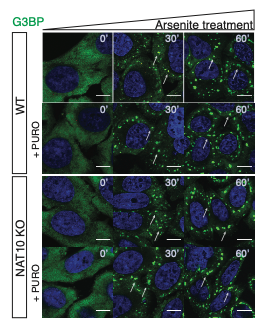

**Supplementary Figure 3. SG formation in response to sodium arsenite treatment**

Microscopy images of G3BP accumulation into SGs in response to treatment with 0.5 mM NaAsO_2_ with or without 10 μg/ml puromycin for 0 min, 30 min or 60 min in HeLa WT and NAT10 KO cells. Scale bar, 50 μm

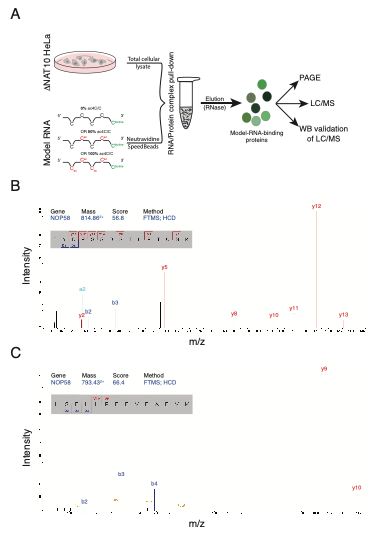

**Supplementary Figure 4. Mass spectrometry identification of NOP58 as an ac4C-binding protein**

The experimental strategy to identify ac4C-binding proteins is shown in panel (a). Panels (b and c) show peptides used to identify NOP58 in mass spectrometry experiments.

**Supplementary Table 1. The SG transcriptome in WT and NAT10 KO HeLa cells**

Overview of average SG enrichment and expression level in rpm of the HeLa transcriptome based on four independent replicates of total RNA and SG RNA from WT and NAT10 KO cells, respectively.

**Supplementary Table 2. Peptide count for proteins identified to bind RNA oligonucleotides**

**Supplementary Table 3. Templates, primers and FISH probes.**
